# Relationships between measures of neurovascular integrity and fluid transport in aging: a multi-modal neuroimaging study

**DOI:** 10.1101/2024.11.05.622194

**Authors:** Ella Rowsthorn, Lachlan Cribb, Benjamin Sinclair, William Pham, Trevor Chong, Stephanie Yiallourou, Marina Cavuoto, Lucy Vivash, Terence J. O’Brien, Xingfeng Shao, Danny J.J. Wang, Meng Law, Matthew P. Pase, Ian H. Harding

## Abstract

Fluid transport in the neurovascular unit (NVU) is essential for maintaining brain health through nutrient delivery and waste clearance. NVU integrity and fluid regulation can be assessed through MRI measures, including water exchange rate through the NVU (BBB k_w_), enlarged perivascular spaces (ePVS), cerebral blood flow (CBF), free water (FW), and white matter hyperintensities (WMH). This study investigated relationships between these MRI measures using Bayesian mixed models, and their variation with chronological age or biological brain age (brainageR) using linear regression in 132 non-clinical older adults (mean age=67 years; 68% female). BBB k_w_ positively associated with CBF (*β*^=0.08, 95% credible interval (CI)=[0.02,0.15]). FW positively associated with both ePVS (*β*^=0.44, CI=[0.30,0.63]) and WMH (*β*^=0.13, CI=[0.04,0.21]). BBB k_w_, CBF and ePVS decreased with age, while FW and WMH increased (all *p*<.05). There were no associations with brain age (all *p*>.05). Relationships between FW, ePVS and WMH likely reflect interconnectivity of fluid regulation within different compartments, while the relationship between BBB k_w_ and CBF indicates a link between NVU fluid flow and vessel function. While individual metrics of NVU integrity are associated with age, their inter-relationships appear stable, providing a baseline for future research in fluid transport and vascular health in neurodegenerative disease.

Graphical Abstract:
Neurovascular function and the fluid transport system.
The brain’s fluid transport system comprises the neurovascular unit, interstitial fluid exchange in the parenchyma and venous outflow. **a.** The flow of water molecules from the vessel to the perivascular space through the blood-brain barrier, and subsequently into the brain (facilitated by aquaporin-4 (AQP4) channels) in quantified by BBB k_w_. **b.** Greater cerebral blood flow rate (CBF) is associated with greater BBB k_w_, possibly due to vessel pulsations increasing osmotic pressure. **c.** Enlarged perivascular spaces (ePVS) may indicate reduced fluid flow within the NVU and are associated with (**d.**) increased extracellular free water (FW) in the parenchyma, suggesting a link between fluid regulation within the vasculature and the surrounding brain tissue. Greater white matter hyperintensity (WMH) volume also occurs alongside increase FW, in line with WMH reflecting more severe fluid flow stagnation occurring in concert with neuroinflammation and/or neuronal demyelination.

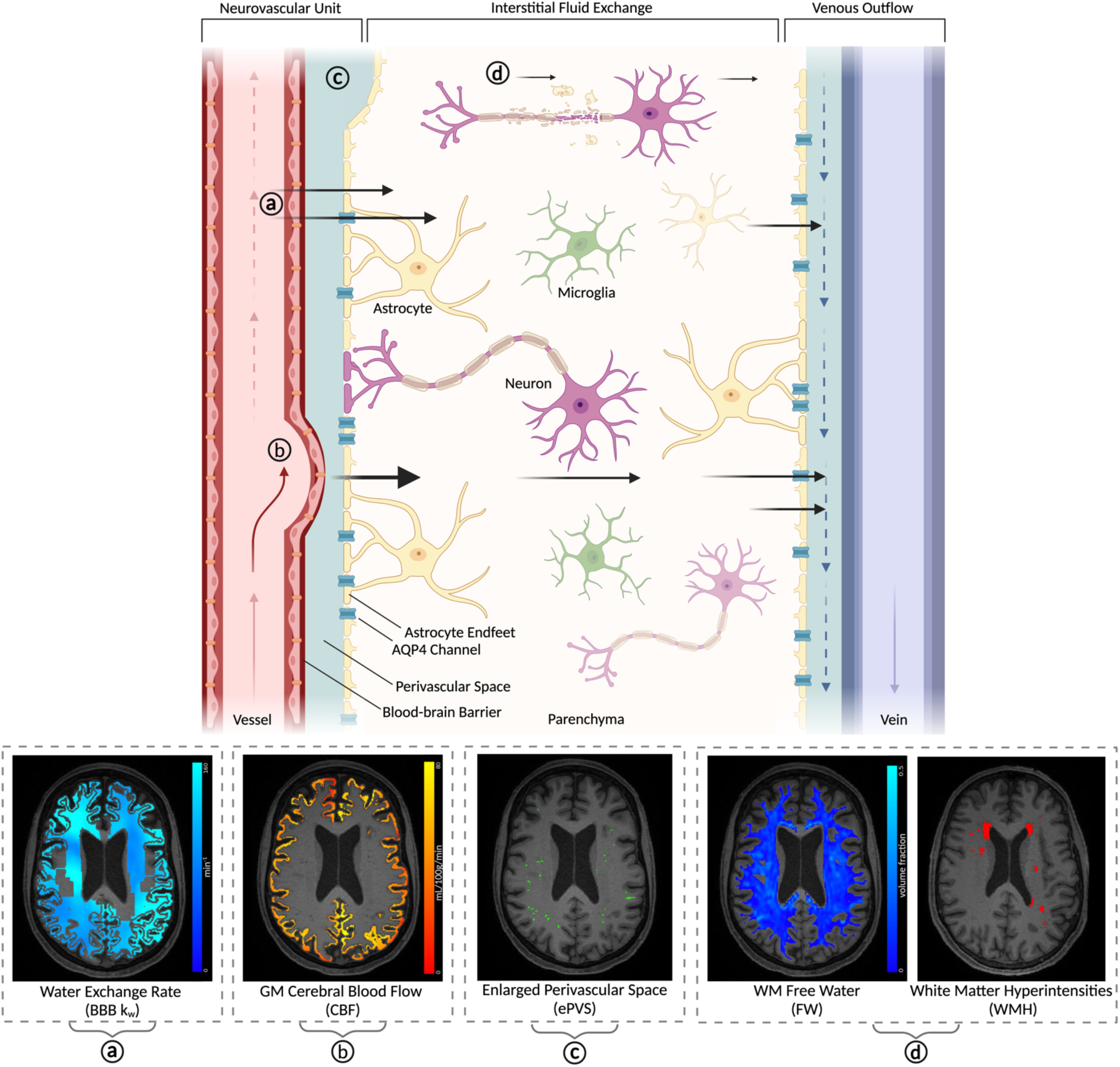

## Introduction

Fluid transport between the neurovasculature and the brain is essential for waste clearance and maintaining homeostasis in the central nervous system. This transport is facilitated by the neurovascular unit (NVU), which comprises several key components including the blood vessel, blood-brain barrier (BBB) formed by endothelial cells and tight junctions, the perivascular spaces that are essential for cerebrospinal fluid (CSF) flow in the brain, and astrocytes that host aquaporin-4 (AQP4) channels which moderate CSF flow into the parenchyma^1,2^. Neuronal, microglial and astrocyte activity influences NVU function, dynamically coordinating fluid regulation in response to neural activity in the local environment^2^. Disruption of fluid regulation, or NVU breakdown, can impair the exchange between CSF and interstitial fluid (ISF) in the parenchyma, which leads to increases in neurotoxic protein accumulation^3–7^. Research has demonstrated that NVU integrity declines with aging, and is further impacted in neurodegenerative disease and cognitive decline^8–12^, highlighting the importance of understanding fluid transport mechanisms within the brain.

Brain magnetic resonance imaging (MRI) can provide insight into the integrity of the NVU’s different compartments and functional elements. First, the rate of cerebral blood flow (CBF inside the blood vessel can be measured by arterial spin labelling (ASL)^13^. CBF is driven by the metabolic demands of local neurons but is limited by the vessel’s structural and physiological capacity to deliver blood efficiently^14^. CBF is negatively associated with age, but is also notably lower in cognitive decline and Alzheimer’s disease^15,16^. Second, MRI-visible enlarged perivascular spaces (ePVS) primarily represent stagnation of fluid flow within the perivascular space of the NVU^17,18^. A larger volume or number of ePVS, is associated with a higher vascular risk factor burden^19^, cerebral small vessel disease (CSVD)^20^ and dementia^21,22^. Third, Free Water (FW) is the volume fraction of unconstrained (freely diffusing) water molecules in the extracellular space of brain tissue, derived from diffusion-weighted imaging (DWI). FW is a general composite measure of extracellular fluid accumulation resulting from white matter degeneration or fluid stagnation in the brain^23,24^, and is associated with vascular dysfunction and neurodegenerative pathology^25,26^. White matter hyperintensities (WMH), seen using fluid attenuated inversion recovery (FLAIR) imaging, may reflect an even greater or advanced state of fluid dysregulation and associated neuropathology^25,27^. WMH are well researched as a marker of neurovascular health and are a feature of CSVD and many neurodegenerative diseases^28–30^. Thus, several well-established MRI tools are available to characterise NVU function and integrity across its different sub-components.

In addition to established measures of vascular integrity, a non-invasive measurement of fluid transport through the NVU has recently become available. Blood-brain barrier water exchange rate (BBB k_w_) assesses fluid transport from the small vessels of the NVU to the brain parenchyma. Using a specialised MRI sequence incorporating both diffusion weighting and ASL, BBB k_w_ quantifies the rate of water molecule flow from within the NVU vessels passing through the perivascular space and AQP4 channels into the brain^31–33^. Early research has shown that lower BBB k_w_ is associated with reduced AQP4 channel expression^34^ and AQP4 inhibition^35^ in animals, demonstrating sensitivity to the integrity of these compartments of the NVU. Although research is nascent, in humans, BBB k_w_ is reduced in those who are older or who have a higher number of vascular risk factors^33,36^.

To date, research has typically investigated individual elements of NVU function and fluid regulation in aging and diseases using a single MRI measure or using multiple measures that are considered independently. However, growing evidence highlights that elements of NVU function and fluid transport likely operate as a complex and highly interdependent system, where the dynamic relationships between components are important for homeostasis^24^. Investigating the relationships *between* elements of the NVU may offer a more comprehensive understanding of NVU function and help guide future research into NVU dysfunction in disease. Accordingly, we aimed to investigate the relationships between MRI measures of NVU integrity, including BBB k_w_, CBF, ePVS, FW and WMH. Additionally, we investigated these measures in association with chronological age and biological brain age^37^. Understanding the relationship between fluid transport and measures of NVU integrity, and how these relationships may change with age, will provide a richer understanding of NVU and fluid transport mechanisms as an integrated system.

## Methods

### Study Participants

MRI data was collected as part of the Brain and Cognitive Health (BACH) cohort study, a prospective cohort study that recruits older adults from the community. To be eligible, participants were 55-80 years of age, without self-reported significant neurological disease (e.g., without dementia, Parkinson’s disease, prior disabling stroke, etc.) and willing to attend in-person assessments at the Alfred Hospital, Melbourne, Australia. Exclusion criteria included contraindication to MRI, and screening failure for moderate-severe cognitive impairment (assessed with the TELE^38,39^). The BACH cohort study was approved by the Alfred Health Ethics Committee (project 78642) and each participant provided written informed consent. This study used data from the first 149 participants, obtained from 2022 to 2023.

### MRI Data Acquisition

All participants were scanned on a 3T Siemens Prisma MRI machine using a 64-channel head coil. The scan session was approximately 45 minutes in duration, and included T1-weighted, T2-weighted FLAIR, diffusion weighted imaging (DWI), pseudo-continuous ASL (pCASL) and diffusion-prepared pCASL (DP-pCASL) acquisitions.

T1-weighted parameters: 3D magnetisation-prepared rapid gradient-echo (MPRAGE) sequence with TR 2400ms, TE 2.24ms, flip angle 8°, FOV 240 x 256 x 154mm, 192 sagittal slices of 0.8mm thickness, resolution 0.8 x 0.8 x 0.8mm.

FLAIR parameters: TR 6700ms, TE 463ms, inversion time 2200ms, variable flip angle, FOV 256 x 256 x 154mm, 192 sagittal slices of 0.8mm thickness, resolution 0.8 x 0.8 x 0.8m.

DWI parameters: Posterior-to-Anterior (P>A) phase direction with TR 3500ms, TE 71.0ms, FOV 232 x 232 x 162mm, 81 axial slices, resolution 2.0 x 2.0 x 2.0mm^3^, 127 total volumes(13 directions at b=0s/mm^2^, 6 directions at b=500s/mm^2^, 48 directions at b=1000s/mm^2^ and 60 directions at b=2000s/mm^2^). An additional echo-planar 2D image with Anterior-to-Posterior (A>P) phase direction with otherwise equivalent acquisition parameters was collected (b-value 0s/mm^2^) for gradient bias correction.

pCASL parameters: TR 4100ms, TE 36.48ms, flip angle 120°, FOV 240 x 240 x 120mm, 40 axial slices, resolution 2.5 x 2.5 x 3mm, label duration = 1500ms, multiple post-labelling delay times and repetitions (500ms x 2 repetitions, 1000ms x 2, 1500ms x 2, 2.000ms x 3, 2500ms x 3), background suppression (except for first M0 reference image with post-labelling delay of 2000ms).

DP-pCASL parameters^33^: TR 4200ms, TE 36.26ms, flip angle 120°, FOV 224 x 224 x 96mm, 12 axial slices, resolution 3.5 x 3.5 x 8mm, label duration = 1500ms, post-labelling delay times, repetitions and b-values (900ms with b=0s/mm^2^ x 15 repetitions, 1800ms with b=0s/mm^2^ x 20, 900ms with b=14s/mm^2^ x 15, 1800ms with b=50s/mm^2^ x 20, 2000ms with b=0s/mm^2^ x 1).

### MRI Processing

#### Intracranial Volume and Structural Segmentations

T1-weighted images were processed through Fastsurfer^40^ (v1.0.0, e4ed6f7) to derive the estimated total intracranial volume (eTIV) and generation of brain segmentations relevant to each MRI outcome (e.g. regional gray matter and white matter), as described further below. All Fastsurfer segmentations were eroded by a 1mm radius spherical kernel to mitigate partial-volume effects at the tissue boundaries.

#### BBB k_w_

BBB k_w_ was derived from the DP-pCASL sequence using the Water Exchange Rate Quantification Toolbox from the Laboratory Of Functional MRI Technology (LOFT), as described previously^33^. Briefly, blood travelling to the brain is tagged within the carotid artery and flows into the small vessels where it is imaged after a sufficient delay via spin-echo ASL. Simultaneous diffusion-weighting allows for water molecules that are flowing along the vessels to be identified separately from the water exchanging into parenchyma through the BBB. The data is first pre-processed to account for head motion and physiological noise, and voxels likely to represent large arteries or ventricles based on a tissue-probability map from SPM were excluded. BBB k_w_ is then estimated using a single-pass approximation model^33^, and quantified as the average rate of water molecules exchanging from capillaries into brain tissue at each voxel in the imaging field of view. The derived BBB k_w_ intensity maps were then manually co-registered to the participant’s native T1w image space using FSLeyes (FSL v6.0.5) and masked to the gray and white matter using segmentations from Fastsurfer.

#### WMH Volume

The volume of WMH was calculated using an in-house deep learning model to segment WMH from FLAIR images. An nnU-Net model^41^ was trained for our specific dataset by first using the SPM lesion segmentation tool (LST) to label WMHs on 47 FLAIR images, and subsequently manually correcting the labels. The trained nnU-Net was then employed to automatically segment WMHs for all participants, and the resulting segmentations were visually inspected for accuracy. Due to potential head movement between T1-weighted and FLAIR acquisitions, FLAIR images were registered to the T1-weighted space via Advanced Normalization Tools^42^ (ANTs, v20190910) to ensure voxel correspondence. Using this registration, WMH segmentations were then transformed to the T1-weighted space using a linear transformation and nearest-neighbour interpolation.

Mean WMH volumes across the whole brain and within regions (see sub-section *Vascular Territory Brain Regions* below) were added to 1 and log transformed, due to a highly skewed distribution and possible values of 0.

#### ePVS Volume Fraction in the White Matter and Deep Gray Matter

The volume of ePVS was quantified using a deep learning model, PINGU, applied to the T1-weighted images. PINGU is a nnU-Net model trained on a heterogenous sample of manually segmented T1-weighted scans for broad-use^43^. We validated PINGU for use in our dataset (**Supplement**). Briefly, a trained rater (ER) completed manual ePVS segmentation of axial slices in a subset of 10 participants, reviewed by a senior neuroradiologist (ML). Automated ePVS segmentations from PINGU were then compared to the manual segmentations. PINGU was in high agreeance, achieving an average correlation of *r* = 0.834 and average Dice score of 0.447 (consistent with previous performance and expectations^43^).

The presence of WMH can confound the identification and quantification of ePVS. Although most prominent on FLAIR images, severe WMH also appear as hypointense areas on T1-weighted images. As such, T1 hypointensities may occlude the detection of ePVS in these areas. Furthermore, it is possible that very small voxel clusters of WMH on FLAIR could be misidentified as ePVS-presenting hypointensities on T1-weighted imaging. Given this, we removed voxels identified as WMH on FLAIR from the PINGU ePVS segmentations. Finally, we quantified ePVS volume fraction as the ratio of volume of ePVS divided by the total normal-appearing white matter and deep gray matter volume (i.e., the caudate, putamen, pallidum, accumbens, thalamus, ventral diencephalon) using segmentations from Fastsurfer.

#### White Matter Free Water (Isotropic Volume Fraction)

Free water (FW) was quantified using the Neurite Orientation Dispersion and Density Imaging (NODDI) toolbox, applied to the DWI data. The NODDI algorithm models water diffusion in each voxel of the images into three fractions: highly restricted intracellular movement, moderately restricted peri-axonal movement and unrestricted isotropic movement (FW)^44^. A higher fraction of FW in a voxel represents a greater proportion of freely diffusing molecules.

To quantify FW, firstly, DWI images were pre-processed through QSIPrep (version 0.14.3, based on Nipype 1.6.1) for denoising, distortion correction, motion correction, normalisation and quality checking. The images were then processed through the NODDI toolbox for Matlab (version 1.0.5, Matlab R2019b)^44^. The DWI b0 was registered to the participant’s T1-weighted image via ANTs, where the whole brain FW intensity maps were then linearly transformed to the T1-weighted space using this registration.

There were substantial spill-over effects of FW estimates near the lateral ventricles (values approaching 1.0), resulting in artificially elevated levels of FW in the neighbouring white matter regions. This likely occurred due to partial volume effects and/or intrinsic spatial smoothness of the diffusion signal. We determined upon careful assessment of the FW intensity maps that masking the data using an upper threshold of 0.5 would sufficiently remove artefactual ventricular spill-over without excluding other white matter voxels (i.e. FW values in the white matter were universally less than 0.5, except at CSF borders; see **eFigure 1** for an example). The resulting FW maps were masked to the white matter using Fastsurfer segmentations.

#### Gray Matter CBF

CBF was derived from pCASL images using the Bayesian Inference for Arterial Spin Labelling MRI (BASIL) toolbox from FSL^45^. The ASL image pairs were first calibrated (standardised) to the M0 image, and then corrected for distortions, motion and partial volume effects. The derived CBF intensity maps were in absolute units (mL/100g/min) and were co-registered to the T1-weighted image within BASIL via FLIRT (v6.0). Output images were masked to the gray matter using the Fastsurfer segmentations.

#### Vascular Territory Brain Regions

To investigate regional effects, we used a previously defined vascular territory atlas^46^. The labels of this atlas were defined as brain regions likely to be perfused by each major artery in each hemisphere, i.e. the anterior, middle or posterior cerebral artery, based on data from over 1200 stroke patients. ANTs was used to register and non-linearly transform the atlas to each subject’s T1w native space. Lateral ventricle segmentations from Fastsurfer were imposed on the resulting atlas transformation to improve ventricle labelling and ensure that the brain atlas regions did not include ventricle voxels. As the cerebellum is not captured by the DP-pCASL sequence, the vertebro-basilar regions of the atlas were not included in regional analyses. The mean values or total volume within the anterior, middle and posterior cerebral artery regions in each hemisphere were quantified for all MRI outcomes using FSL. An example of the final resulting atlas registration is presented in **Figure 1**.

**Figure 1.**
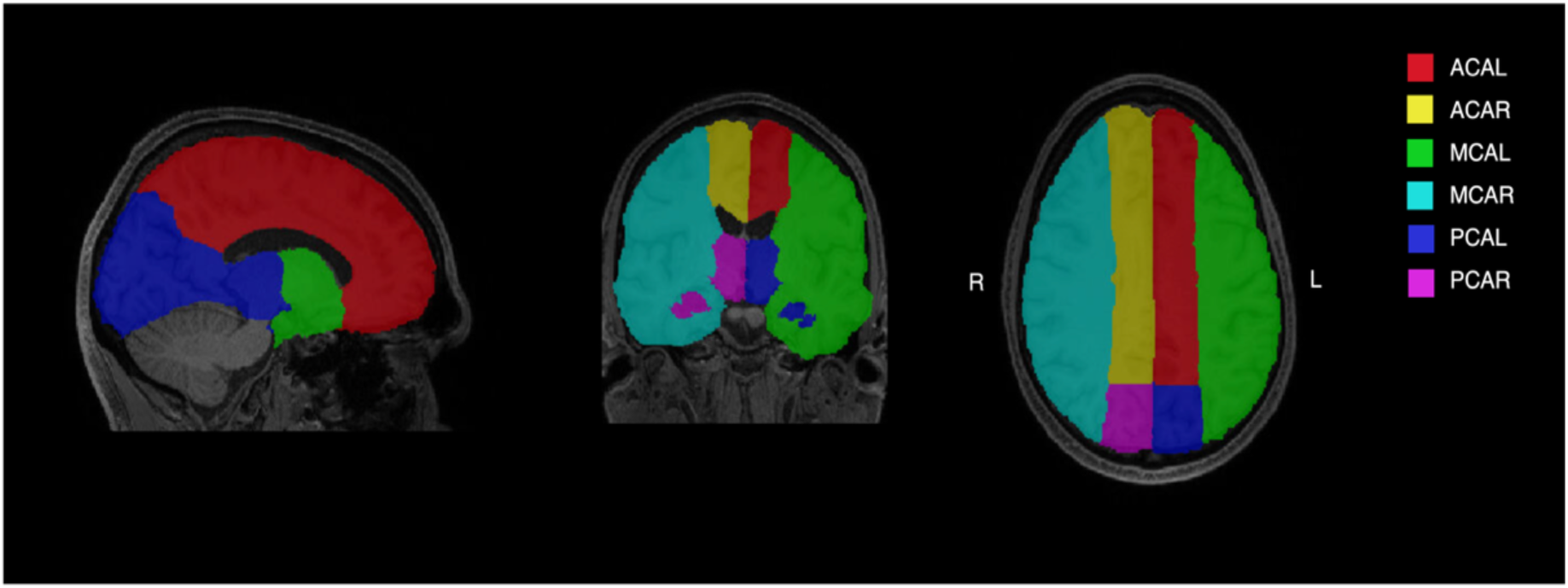
The vascular territory atlas registered to a T1w image. Atlas labels were defined as brain regions likely to be perfused by each major artery in each hemisphere^46^. ACAL = left anterior cerebral artery region, ACAR = right anterior cerebral artery region, MCAL = left middle cerebral artery region, MCAR = right middle cerebral artery region, PCAL = left posterior cerebral artery region, PCAR = right posterior cerebral artery region.

#### Brainage (brainPAD)

Volumetric brain age was derived from brainageR, as described elsewhere^37,47^. Briefly, brainageR pre-processes T1w images through SPM12 to generate voxel-level gray and white matter volume estimates across the whole brain. Then, a previously trained brainage estimate model in R (via kernlab^48^) is used to predict brain age using the gray and white matter volume vectors. We then calculated the difference between the predicted brain age and chronological age to derive the brain predicted age difference (brainPAD). Positive brainPAD values indicate an older estimated age relative to true age.

### Data Analysis

To evaluate the global associations between measures of the NVU and to assess whether these associations varied by region, a Bayesian mixed effects model was fitted. Bayesian modelling was used because: 1) we expected MRI outcomes to be substantially correlated within subjects (e.g., a participant with high values of BBB k_w_ in a one region would likely have high values in other regions) – Bayesian modelling manages this expectation through shrinkage^49^; 2) it allowed the estimation of a large number of model parameters, capturing variation in the relationship between MRI outcomes across subjects and regions, relative to the available sample size; and 3) it provides exact inferences and reduced mean squared error even in smaller samples.

Using the *brms* package^50^ in R, the following Bayesian model was fitted for each pair of MRI outcome variables (e.g. BBB k_w_ and FW):

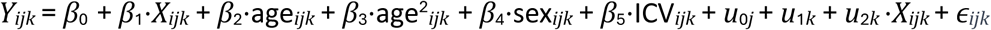

*Y* and *X* represent two of the MRI variables as outcome and predictors respectively. *i* indexes observation within *j* subject and *k* indexes the six atlas region levels. The random intercept for subject is represented by *u*_0_*_j_*, the random intercept for region is represented by *u*_1_*_k_* and the random slope of *X* across regions is represented by *u*_2_*_k_* ⋅*X_ijk_*. Sex, eTIV and second-order polynomial age were included as covariates. We estimated each model using Markov Chain Monte Carlo (MCMC) sampling with 4 chains of 5500 iterations. Priors over parameters were set as exponential for all standard deviations and sigma (*e*^1/*x*^), Lewandowski-Kurowicka-Joe (LKJ) Cholesky factor distributions for the correlation matrix (eta=2) and normal Gaussian distributions for the global intercept and remaining model terms (location = 0, scale = 2). R-hat values below 1.01 were considered to represent stability and likely convergence^51^, and effective sample sizes (ESS) greater than 1000 were considered to reflect reliability^50^. 95% credible intervals (CI; highest density interval) that do not overlap the null provide evidence of a given relationship, where the effect size is reflected by the standardised median posterior distribution estimate (*β*^).

Linear regression models were used to investigate the global associations between each MRI outcome, chronological age and brainPAD. As some of the MRI outcomes have previously been found to have non-linear relationships with age^52^, an additional quadratic term for age was included in all models. To determine whether quadratic age better explained variance in MRI outcomes than linear age, Akaike Information Criterion (AIC) was used to compare models with linear or second-order polynomial age terms. Furthermore, to explore whether age modified the global-level relationships *between* MRI outcomes, linear regressions with an interaction term for age were also tested. This was repeated using an interaction term for brainPAD. All models were adjusted for sex, intracranial volume and age. A p-value < .05 was considered statistically significant.

## Results

After exclusion of incomplete (n=6) or poor-quality (n=11) MRI scans, 132 participants had complete MRI data available for analysis. The mean age was 66.95 years (*SD* = 5.32) and 68.2% of the sample were women. Mean brainPAD was −4.26 years (*SD* = 5.87), meaning that on average, participants demonstrated a brain age slightly younger than their chronological age (**Table 1**).

**Table 1.** Cohort Summary.

|  | Sample<br>(N=132) |
| --- | --- |
| Women (n, %) | 90 (68.2%) |
| Age, years | 66.95 ± 5.32 |
| Brainage, years | 62.69 ± 7.79 |
| brainPAD, years | -4.26 ± 5.87 |
| BBB Water Exchange Rate, min<sup>-1</sup> | 117.08 ± 24.48 |
| Free Water, volume fraction | 0.07 ± 0.01 |
| ePVS volume, mm<sup>3</sup> (median, IQR) | 2712.06 ± 2010.11 |
| ePVS, volume fraction | 0.006 ± 0.003 |
| Cerebral Blood Flow, mL/100g/min | 38.31 ± 8.32 |
| WMH volume, mm<sup>3</sup> (median, IQR) | 532.74 ± 1022.34 |
| WMH volume, log +1 | 6.32 ± 1.54 |

**Note:** Reported as mean ± standard deviation unless otherwise specified. MRI outcome measures reflect whole brain means within the grey matter, white matter, or both, as described further in the text.
BBB = blood-brain barrier, brainPAD = volumetric brain age minus chronological age; ePVS = enlarged perivascular space.

### Associations between elements of the NVU

Relationships between all MRI outcome pairs are summarised in **Table 2**, and in reported in detail in the **Supplement**. A positive global relationship was evident between BBB k_w_ and CBF *β*^ = 0.15, 95% CI = [0.07, 0.22]) (**Figure 2a**). There was also a positive global relationship between FW and ePVS (*β*^ = 0.44, 95% CI = [0.30, 0.63]; **Figure 2b**), and a positive global relationship between FW and WMH (*β*^ = 0.13, 95% CI = [0.04, 0.21]; **Figure 2c**). There was negligible evidence to suggest any relationships between the other MRI outcome pairs (**eTable 2 – eTable 11**).

**Figure 2.**
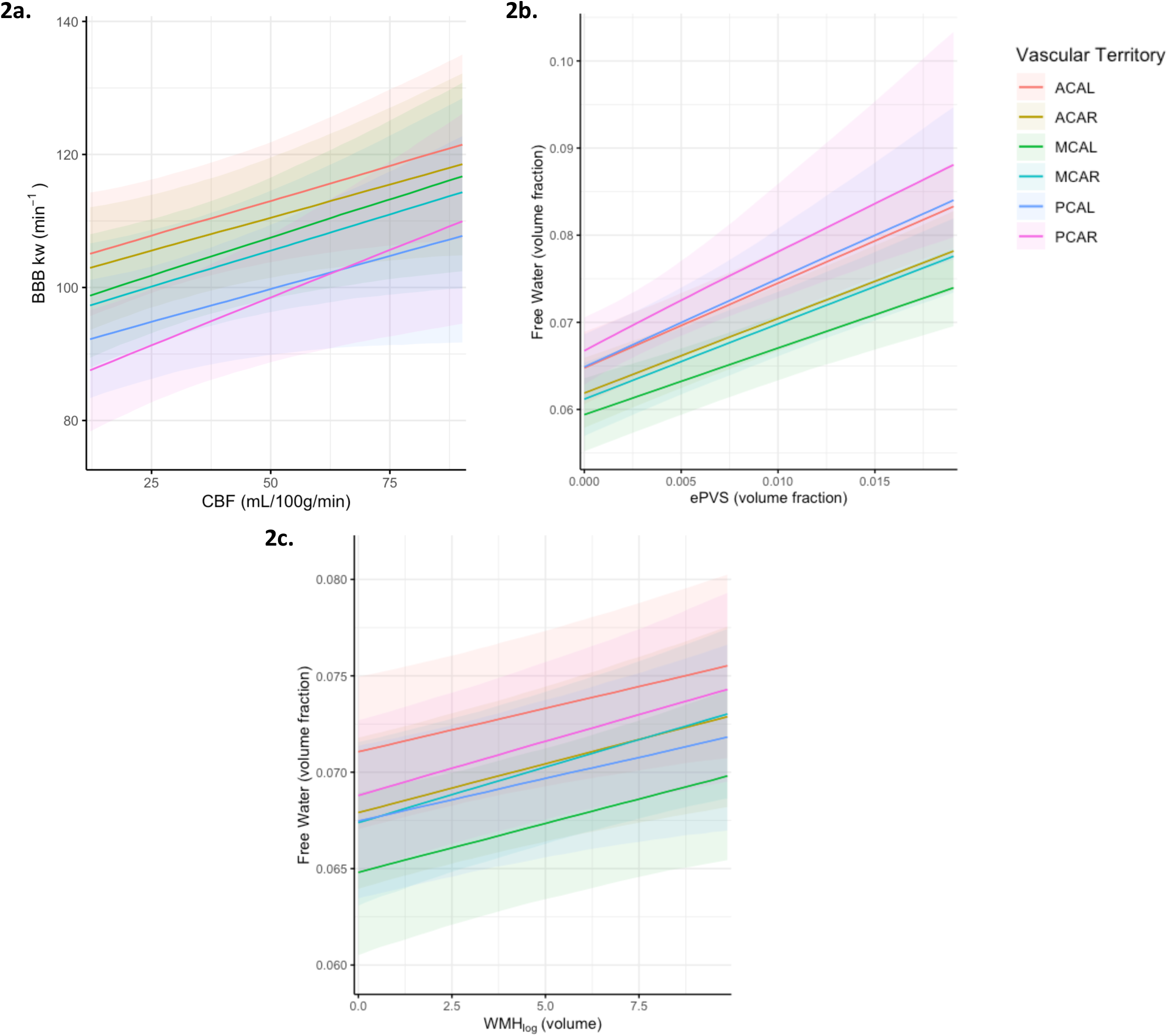
Associations between MRI measures in all vascular atlas territories. All plotted relationships show evidence in favour of an overall association between the measures (95% credible interval that does not cross zero) with no observed difference across vascular territory, indicating a consistent relationship throughout the brain. BBB k_w_ = blood-brain barrier water exchange rate; CBF = gray matter cerebral blood flow; ePVS = enlarged perivascular space volume fraction; FW = Free Water volume fraction; WMH_log_ = log transformed white matter hyperintensity volume. ACAL = left anterior cerebral artery region, ACAR = right anterior cerebral artery region, MCAL = left middle cerebral artery region, MCAR = right middle cerebral artery region, PCAL = left posterior cerebral artery region, PCAR = right posterior cerebral artery region.

**Table 2.** Relationships between elements of the NVU.

| | BBB $k_w$ | | | FW | | | ePVS | | | CBF | | |
| --- | --- | --- | --- | --- | --- | --- | --- | --- | --- | --- | --- | --- |
| | $\beta^{\wedge}$ | 95% CI | $R^2_{\text{marginal}}$ | $\beta^{\wedge}$ | 95% CI | $R^2_{\text{marginal}}$ | $\beta^{\wedge}$ | 95% CI | $R^2_{\text{marginal}}$ | $\beta^{\wedge}$ | 95% CI | $R^2_{\text{marginal}}$ |
| FW | -0.04 | -0.10, 0.02 | 0.14 | - |  |  | - |  |  | - |  |  |
| ePVS | 0.03 | -0.06, 0.14 | 0.12 | **0.44** | **0.30, 0.63** | **0.26** | - |  |  | - |  |  |
| CBF | **0.08** | **0.02, 0.15** | **0.15** | 0.07 | -0.01, 0.15 | 0.13 | 0.01 | -0.06, 0.08 | 0.07 | - |  |  |
| WMH | 0.01 | -0.06, 0.07 | 0.14 | **0.13** | **0.04, 0.21** | **0.13** | 0.06 | -0.03, 0.14 | 0.08 | -0.00 | -0.07, 0.07 | 0.10 |

**Note:** Column = outcome variable. Row = predictor variable.
$\beta^{\wedge}$ = posterior distribution estimate; 95% CI = 95% credible interval; $R^2_{\text{marginal}}$ = proportion of variance explained by fixed effects; bolding indicates evidence against the null.
BBB $k_w$ = blood-brain barrier water exchange rate; CBF = gray matter cerebral blood flow; ePVS = enlarged perivascular space volume fraction;
FW = Free Water volume fraction; WMH = log transformed white matter hyperintensity volume.

### Associations between elements of the NVU, Chronological Age and Brainage

BBB k_w,_ CBF and ePVS significantly decreased with age, while FW and WMH volume significantly increased with age (all *p*<.05; **Figure 3; eTable12**). Models fitted with either linear or quadratic age terms were comparable for all measures (AIC difference ≤2 for all). Plots with a fitted quadratic line can be found in the supplement (**eFigure2**).

**Figure 3.**
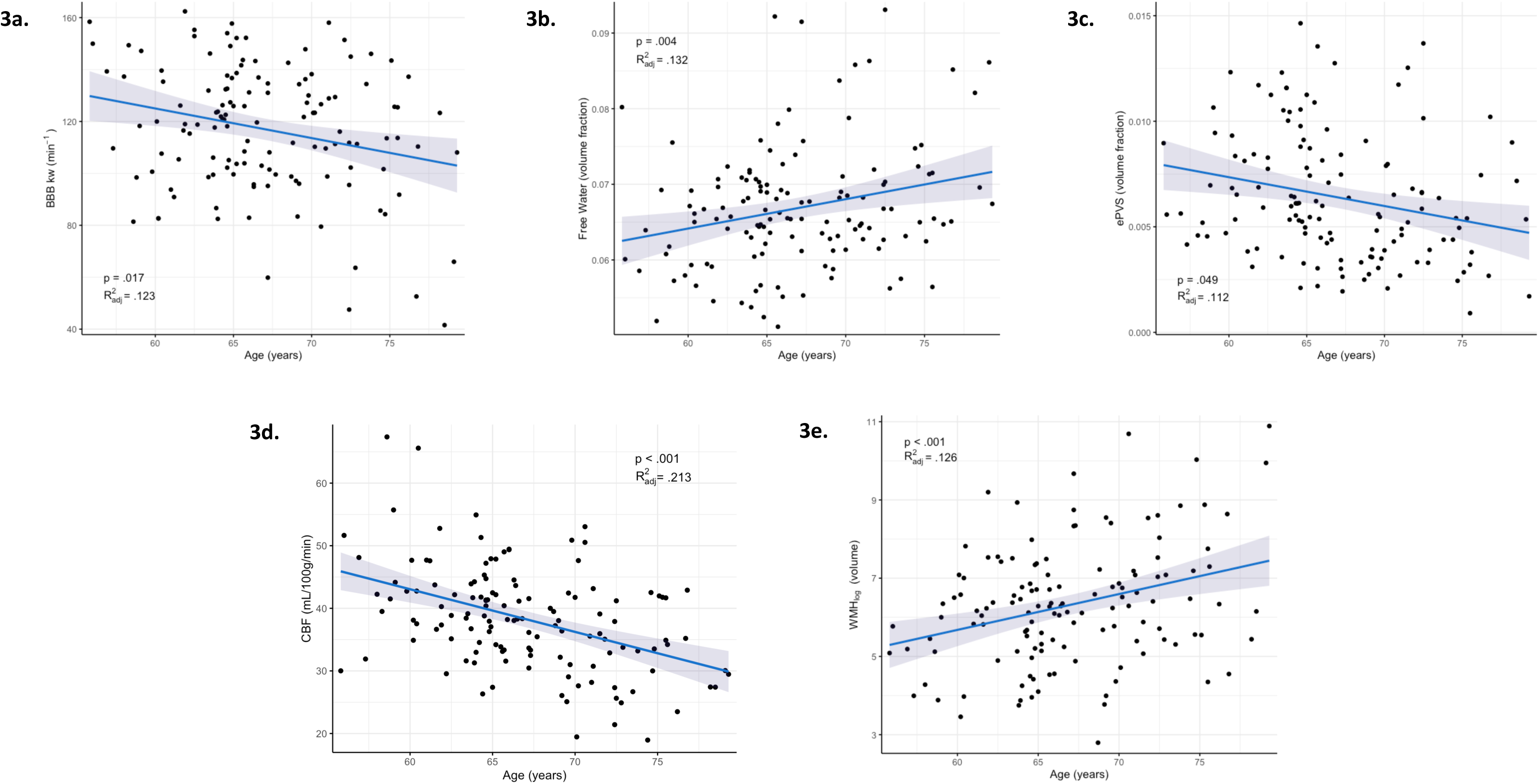
Associations between each MRI outcome (whole brain average) and age. All relationships are statistically significant (*p*<.05). BBB k_w_ = blood-brain barrier water exchange rate; CBF = gray matter cerebral blood flow; enlarged perivascular space volume fraction; FW = Free Water volume fraction; WMH_log_ = log transformed white matter hyperintensity volume.

The relationship between ePVS volume fraction and WMH volume was modified by age and age^2^ (*p*=.029). To further explore this non-linear effect modification beyond a quadratic fit, we conducted repeated this regression analysis with a spline term (basis spline with four degrees of freedom) for age in a post-hoc analysis. We found that in older participants, WMH volume is elevated in those with lower ePVS but decreased in those with higher ePVS volumes (p=.012; **Figure 4**). Age did not significantly modify the relationship between other MRI outcome pairs (all p>.05).

**Figure 4.**
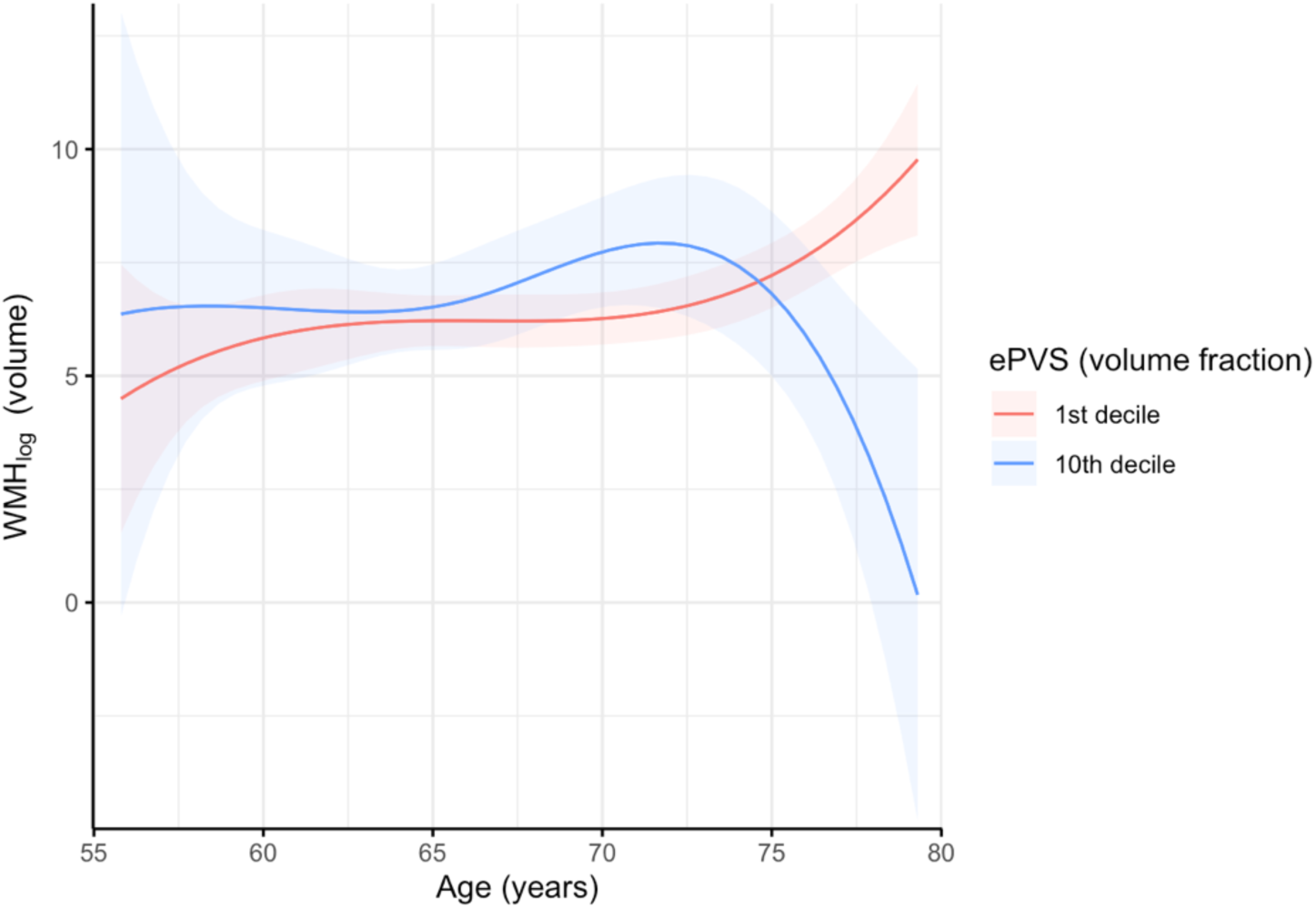
Age modifies the relationship between ePVS and WMH. Linear regression between ePVS volume fraction and WMH volume with an interaction spline term for age. ePVS volume is plotted at the 1^st^ and 10^th^ decile of the sample. ePVS = enlarged perivascular space volume fraction; WMH_log_ = log transformed white matter hyperintensity volume.

brainPAD was not associated with any of the MRI outcomes (all *p*>.05) and did not significantly modify the relationship between any MRI outcome pairs (*p*>.05; **eTable 14**).

## Discussion

This study aimed to investigate the associations between functional elements of the NVU, and age in a community cohort. We found that more fluid accumulation within the NVU (ePVS) was associated with a greater quantity of extracellular fluid in the parenchyma (FW), which was in turn associated with more profuse signatures of vascular-associated neuropathology (WMH). As such, these measures may reflect dysfunction or stagnation of fluid flow within different compartments of an interconnected fluid transport system. We also found that greater blood flow through the vessel (CBF) was associated with faster fluid transport through the perivascular spaces and astrocyte endfeet (BBB k_w_), suggesting that vessel integrity or activity may have an important role in driving fluid flow within the NVU. Individually, most markers of NVU integrity showed putative worsening with age, but the inter-relationships between these measures were largely invariant to age. Together, our findings highlight inter-dependencies between elements of neurovascular function which provide important insight into NVU as an integrated system.

### An Interconnected Fluid Transport System

In the brain, the fluid transport system is predominantly formed by three compartments: the perivascular space in the NVU that facilitates transfer of CSF into the brain; the CSF-ISF exchange in the extracellular space of the brain parenchyma; and the venous ‘perivascular space’ that transports CSF out of the brain^53^ (although further clarification of venous transport mechanisms is needed^54^). We found evidence supporting inter-dependencies between these compartments, where enlargement of the perivascular space was related to fluid accumulation in the extracellular space where CSF-ISF exchange occurs. This link between CSF fluid flow within the vasculature and brain tissue itself is consistent with previous research^24,55^, and suggests that both measures may reflect the fluid stagnation within different compartments of a highly inter-dependent fluid transport system. Although we cannot determine causality within this cross-sectional study, animal research and multi-compartmental modelling studies propose that this relationship is likely bi-directional^56,57^.

We also found a relationship between FW and WMH, which is in line with previous research suggesting that WMH formation is preceded by an increase in local FW^27^ and is a more severe or later-stage marker of fluid accumulation (in concert with neuropathology^58,59^). Although we didn’t find a direct relationship between ePVS and WMH, we believe this is due to methodological limitations (see section *Study Strengths and Limitations*), rather than a lack of biological association. Further research into the associations between additional indicators of fluid transport, such as diffusion along the perivascular space (DTI-ALPS)^60,61^, or CSF biomarkers of protein aggregate clearance (e.g. beta-amyloid and tau), is needed to continue to elucidate the dependencies and consequences of fluid flow stagnation within this system.

Interestingly, we did not replicate a previous finding where lower BBB k_w_, reflecting fluid transport through the NVU, was associated with increased free water in the parenchyma^62^. Given that fluid flow through this system is largely dependent on net osmotic pressure and ionic concentration gradients present throughout the whole network^63^, it is plausible that reduced flow through the NVU would coincide with reduced flow in the CSF-ISF exchange compartment. Further research is needed to determine whether this relationship perhaps only emerges in specific contexts or with more severe impairment to the fluid transport system.

### Vessel Function Supports Fluid Transport in the NVU

Flow through the fluid transport system is partly driven by the volume changes resulting from pulsatile vasodilation of the vessel in the NVU^64–66^. During vasodilation, the vessel expands within the perivascular space, reducing the occupiable area for CSF and thus causing hydrostatic pressure that encourages flow through astrocyte end-feet and AQP4 channels^67^. Concordantly, vasodilation and vasoconstriction in response to neural activity, termed neurovascular coupling, has been shown to modulate flow through the waste clearance network^68^, especially during sleep^69,70^. Taken together, this research suggests that blood vessel function likely affects the capacity for fluid flow both within the NVU and, in turn, through the fluid transport system. In our study, BBB k_w_ was positively associated with CBF, in concordance with previous research reporting that BBB k_w_ values seem to converge with CBF^71^. Given that CBF represents the net momentum of vasodilation and vasoconstriction, the association between CBF and BBB k_w_ likely reflects this underlying physiological relationship. This adds to a fast growing body of multi-disciplinary research demonstrating that the vessel in the NVU has an important role in facilitating flow within the fluid transport system^72^.

### Age, Brainage and NVU Integrity

We found that all measures of the NVU changed with age as expected. However, relationships between aspects of neurovascular function or fluid transport were not modified by age, unless explained by methodological limitations (discussed further below). This suggests that the dependencies between elements of the may NVU exist regardless of age-related vascular breakdown, and that alterations to relationships between MRI measures may provide a putative indicator of pathology independent of normal aging.

Although testing this hypothesis is beyond the scope of the current study, this notion has been proposed in previous work, where some inter-relationships seem to only exist in either ‘healthy’ community or disease cohorts^24^. Future research is encouraged to explore this hypothesis within other age groups and in individuals with neurodegenerative pathology.

In the present study, biological brain age relative to true age (brainPAD) was not related to any elements of NVU function or integrity. BrainPAD is thought to be sensitive marker of general brain health and predictor of cognitive status and disease risk^47^. However, the current result may be unsurprising given that our included MRI measures are thought to represent subtle changes related to neurovascular health that can occur prior to changes in cortical thickness^73^ or atrophy^74,75^ which are used in the calculation of brainPAD. Future research is required to investigate whether relationships between these MRI measures and brainPAD emerge temporally, or perhaps in those with more prevalent atrophy.

### Study Strengths and Limitations

This study has several key strengths. Our study incorporates several state-of-the-art sequences which allowed targeted assessments of outcome measures. To our knowledge, this is the first study to utilise these measures in a multi-modal approach to comprehensively assess NVU dependencies. The focus on a relatively healthy, older-age population provides a valuable baseline for understanding NVU function, establishing a reference point for future research into neurodegenerative diseases.

This study is not without limitations. We have interpreted the MRI measures in this study as relatively stable representations of typical vascular function or integrity; however, it is possible that these measures are affected by acute effects such as neuroinflammatory response^76,77^, diet, alcohol consumption and physical activity^78^. Similarly, although not within the scope of the present study, it is possible that the functional relationships elements of the NVU change in the presence of vascular-related conditions such as diabetes^79,80^ or hypertension^81^. While our results provide a general characterisation of the dependencies within the NVU, further work is necessary to investigate the potential influence of these acute factors and to further elucidate the dynamic nature of NVU function.

We also encountered a methodological limitation when exploring the relationship between fluid stagnation within the perivascular space (ePVS) and severe fluid accumulation reflected by WMH. Despite both FW and WMH reflecting fluid accumulation within the parenchyma, we only found an association between ePVS and FW and not between ePVS and WMH. This contrasts with previous research, which has repeatedly reported a positive relationship between visually-graded ePVS and WMH^24^. Methodological differences between the volumetric ePVS approach used herein and previous grading approaches might explain this discrepancy. In particular, we observed that the presence of prominent WMH may occlude MRI-visible ePVS in overlapping areas (even on T1-weighted MRI). As such, it is necessary to quantify ePVS volume as a fraction of normal-appearing white matter.

However, it remains biologically plausible that ePVS are also present – and potentially even predominate – in WMH regions. As a result, the association between ePVS and WMH may falsely weaken as WMH volume increases. This phenomenon might contribute to the age interaction between ePVS and WMH observed in our study. Although both ePVS and WMH volumes independently increase with age^52,82^, having high volumes of both simultaneously is less likely due to their contradictory nature, and achieving this would require overexpression of ePVS in non-WMH areas. Given that WMH volume tends to increase with age, this effect is likely exaggerated in those who are older, leading to the apparent decoupling of the relationship between ePVS and WMH observed in our results. This phenomenon may also explain the unexpected negative trend between brainPAD and ePVS, as older brainage is also associated with greater WMH^83^. Future research should further investigate the WMH-occlusion effect on the detection of ePVS on T1-weighted MRI. In the context of aging and neurodegenerative disease, ePVS may have the most utility as an early indicator of NVU integrity before WMH develops, or instead might require non-linear adjustment in the presence of extensive WMH.

## Conclusion

Associations between elements of NVU function and fluid transport may reflect underlying physiological dependencies between the fluid transport system or within neurovascular components in the brain. Increased extracellular fluid accumulation in the parenchyma (FW) was associated with both perivascular space enlargement (ePVS) and white matter hyperintensities (WMH), likely representative of a highly interconnected fluid transport system. Furthermore, vessel function may help facilitate fluid flow within the NVU, represented by the relationship between cerebral blood flow (CBF) and fluid transport through the NVU (BBB k_w_). Together, the age-independent associations between elements of the NVU found within this non-clinical community cohort provide a foundation for future investigations of neurodegenerative disease progression.

