## Supplement for "Relationships between measures of neurovascular integrity and fluid transport in aging: a multi-modal neuroimaging study"

### Table of Contents


| | |
|---|---|
| Validation of PINGU for use in the present dataset | 1 |
| Free Water thresholding to remove artefactual voxels | 3 |
| Tables of results for relationships between MRI measure pairs | 4 |
| Plotted relationships between MRI measure pairs | 14 |
| Table of results for relationships between MRI measures and age | 15 |
| Table of results for age interactions | 16 |
| Table of results for relationships between MRI measures and brain age | 19 |
| Plotted relationships between quadratic age and MRI measures | 20 |
| References | 21 |

### Validation of PINGU for use in the Present Dataset

From the 132 available participants, 10 were randomly selected for validation of PINGU[1]. Single axial slices at the level of the basal ganglia and in the centrum semiovale white matter were selected in each participant according to proposed advances to STRIVE criteria[2]: the basal ganglia slice was identified as the slice showing the anterior commissure; the white matter slice was identified at 1cm above the lateral ventricles. Voxels of perivascular spaces within slices were labelled by a trained rater (ER) in ITK-Snap (ver 3.6.0) with reference to FLAIR images to ensure voxels of white matter hyperintensities were not labelled. Labels were reviewed by a senior neuroradiologist (ML).

Extracting the same slices, perivascular space segmentations from PINGU were compared to manual segmentations. The mean Dice score of each region was calculated as two times the number of overlapping voxels (i.e. voxels segmented by both methods) divided by the sum of manual and automatically segmented voxels, where a higher Dice score indicates higher similarity (range 0-1). Sensitivity was calculated as the number of overlapping voxels divided by the number of manually segmented voxels, and specificity was calculated as the number of overlapping voxels divided by the number of automatically segmented voxels. The Pearson's correlation between the number of identified voxels within each region was calculated.

Across the 10 participants, the mean Dice score was 0.474 in basal ganglia and 0.447 in the white matter, and the overall Dice score was 0.460 (**eTable 1**). The correlation of the number of identified voxels between PINGU and the manual segmentations was high, with a correlation of $r$ = 0.881 in the basal ganglia, $r$ = 0.787 in the white matter and $r$ = 0.834 overall.

| | Sensitivity | | | Specificity | | | Dice Score | | |
|---|---|---|---|---|---|---|---|---|---|
| **Subject** | BG | WM | Total | BG | WM | Total | BG | WM | Total |
| 1 | 0.257 | 0.197 | 0.227 | 0.925 | 0.833 | 0.879 | 0.403 | 0.319 | 0.361 |
| 2 | 0.390 | 0.560 | 0.475 | 0.695 | 0.699 | 0.697 | 0.500 | 0.622 | 0.561 |
| 3 | 0.115 | 0.463 | 0.289 | 0.206 | 0.506 | 0.356 | 0.148 | 0.484 | 0.316 |
| 4 | 0.341 | 0.584 | 0.462 | 0.547 | 0.500 | 0.524 | 0.420 | 0.539 | 0.479 |
| 5 | 0.741 | 0.923 | 0.832 | 0.661 | 0.187 | 0.424 | 0.699 | 0.311 | 0.505 |
| 6 | 0.397 | 0.380 | 0.388 | 0.828 | 0.857 | 0.843 | 0.537 | 0.527 | 0.532 |
| 7 | 0.287 | 0.100 | 0.193 | 0.807 | 0.700 | 0.754 | 0.424 | 0.175 | 0.299 |
| 8 | 0.373 | 0.360 | 0.366 | 0.528 | 0.580 | 0.554 | 0.437 | 0.444 | 0.44 |
| 9 | 0.500 | 0.272 | 0.386 | 0.715 | 0.409 | 0.562 | 0.588 | 0.327 | 0.457 |
| 10 | 0.433 | 0.779 | 0.606 | 0.878 | 0.676 | 0.777 | 0.580 | 0.724 | 0.652 |
| **Average** | 0.383 | 0.462 | **0.422** | 0.679 | 0.595 | **0.637** | 0.476 | 0.447 | **0.460** |

**eTable 1.** Manual and automatic segmentations of enlarged perivascular spaces.
PINGU perivascular space segmentations were compared to manual segmentations in the basal ganglia and white matter for 10 randomly selected participants. The sensitivity (proportion of true negative voxels), specificity (proportion of true positive voxels) and Dice scores for each participant were calculated in both regions and as a total. Comparisons for all participants are reported.

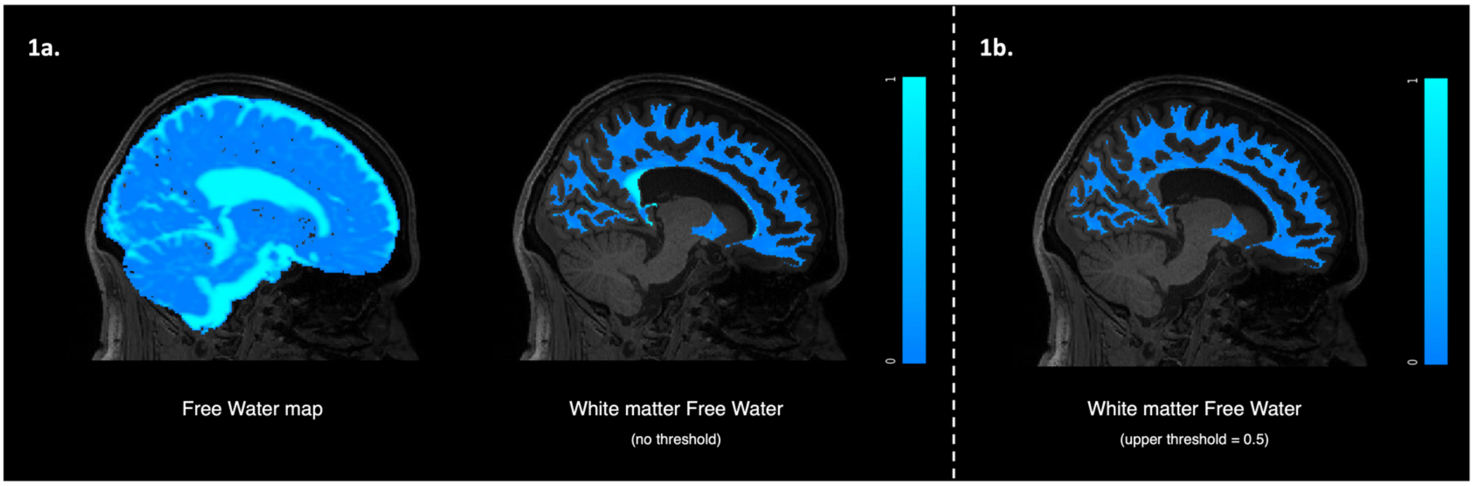


**eFigure 1. Example of Free Water thresholding to remove artefactual ventricular spill-over.**
**1a)** A Free Water map that have been registered to T1w space and then masked to a Fastsurfer-derived white matter segmentation. No thresholding has been applied, and ventricle spill-over is visible (bright blue).
**1b)** The same Free Water map with an upper threshold of 0.5 applied, masked to the white matter segmentation. The ventricle spill-over is no longer present and artefactual voxels have been removed.

**eTable 2 – Relationship between BBB $k_w$ and Free Water across regions.**

| Parameter | Estimate (Error) | 95% CI (low, upp) | R-hat | ESS | Model Fit |
|---|---|---|---|---|---|
| Intercept | 0.29 (0.18) | -0.07, 0.65 | 1.00 | 952 | |
| FW | -0.04 (0.03) | -0.10, 0.02 | 1.00 | 4072 | |
| Age | -0.17 (0.08) | -0.31, -0.01 | 1.00 | 666 | |
| Age^2 | -0.08 (0.06) | -0.20, 0.04 | 1.00 | 673 | |
| Sex (Male) | -0.66 (0.19) | -1.01, -0.29 | 1.00 | 629 | |
| eTIV | 0.12 (0.08) | -0.04, 0.28 | 1.00 | 715 | |
| *sigma* | 0.38 (0.01) | 0.36, 0.40 | 1.00 | 8894 | |
| *sd* of Subject effect (Intercept) | 0.84 (0.05) | 0.74, 0.95 | 1.00 | 1123 | |
| *sd* of Region effect (Intercept) | 0.34 (0.15) | 0.17, 0.72 | 1.00 | 2572 | |
| *sd* of Region effect (FW slope) | 0.03 (0.02) | 0.00, 0.09 | 1.00 | 4242 | |
| *cor* of Region effect (Intercept, FW slope) | -0.11 (0.43) | -0.85, 0.73 | 1.00 | 7465 | |
| $R^2$ | | | | | 0.86 |
| Adjusted $R^2$ | | | | | 0.83 |
| Marginal $R^2$ | | | | | 0.14 |

**Note:** BBB $k_w$ = blood-brain barrier water exchange rate, FW = Free Water volume fraction, eTIV = estimated total intracranial volume. 95% CI = 95 % credible interval**,** *cor* = correlation between the Intercept of region and slope of FW, *sd* = standard deviation of distribution, *sigma* = residual standard deviation.

**eTable 3 – Relationship between BBB $k_w$ and ePVS across regions.**

| Parameter | Estimate (Error) | 95% CI (low, upp) | R-hat | ESS | Model Fit |
|---|---|---|---|---|---|
| Intercept | 0.30 (0.17) | -0.04, 0.64 | 1.00 | 1376 | |
| ePVS | 0.03 (0.05) | -0.06, 0.14 | 1.00 | 3255 | |
| Age | -0.17 (0.08) | -0.33, -0.02 | 1.00 | 926 | |
| Age^2 | -0.08 (0.06) | -0.21, 0.04 | 1.00 | 1064 | |
| Sex (Male) | -0.64 (0.18) | -1.00, -0.29 | 1.00 | 979 | |
| eTIV | 0.11 (0.08) | -0.05, 0.28 | 1.00 | 935 | |
| *sigma* | 0.38 (0.01) | 0.36, 0.40 | 1.00 | 7764 | |
| *sd* of Subject effect (Intercept) | 0.84 (0.05) | 0.74, 0.95 | 1.00 | 1320 | |
| *sd* of Region effect (Intercept) | 0.30 (0.14) | 0.14, 0.66 | 1.00 | 2382 | |
| *sd* of Region effect (ePVS slope) | 0.05 (0.05) | 0.00, 0.17 | 1.00 | 2920 | |
| *cor* of Region effect (Intercept, ePVS slope) | -0.29 (0.43) | -0.92, 0.64 | 1.00 | 6728 | |
| $R^2$ | | | | | 0.85 |
| Adjusted $R^2$ | | | | | 0.82 |
| Marginal $R^2$ | | | | | 0.15 |

**Note:** BBB $k_w$ = blood-brain barrier water exchange rate, ePVS = enlarged perivascular space volume fraction, eTIV = estimated total intracranial volume. 95% CI = 95 % credible interval**,** *cor* = correlation between the Intercept of region and slope of FW, *sd* = standard deviation of distribution, *sigma* = residual standard deviation.

**eTable 4 – Relationship between BBB $k_w$ and CBF across regions.**

| Parameter | Estimate (Error) | 95% CI (low, upp) | R-hat | ESS | Model Fit |
|---|---|---|---|---|---|
| Intercept | 0.26 (0.18) | -0.08, 0.61 | 1.00 | 1346 | |
| CBF | 0.08 (0.04) | 0.02, 0.15 | 1.00 | 3806 | |
| Age | -0.16 (0.08) | -0.31, -0.01 | 1.00 | 1089 | |
| Age^2 | -0.07 (0.06) | -0.19, 0.05 | 1.00 | 1466 | |
| Sex (Male) | -0.62 (0.18) | -0.96, -0.27 | 1.00 | 992 | |
| eTIV | 0.11 (0.08) | -0.05, 0.27 | 1.00 | 1051 | |
| *sigma* | 0.38 (0.01) | 0.36, 0.40 | 1.00 | 8330 | |
| *sd* of Subject effect (Intercept) | 0.84 (0.06) | 0.74, 0.96 | 1.00 | 1264 | |
| *sd* of Region effect (Intercept) | 0.31 (0.14) | 0.15, 0.66 | 1.00 | 2979 | |
| *sd* of Region effect (CBF slope) | 0.03 (0.03) | 0.00, 0.10 | 1.00 | 3870 | |
| *cor* of Region effect (Intercept, CBF slope) | -0.15 (0.43) | -0.85, 0.70 | 1.00 | 8403 | |
| $R^2$ | | | | | 0.86 |
| Adjusted $R^2$ | | | | | 0.83 |
| Marginal $R^2$ | | | | | 0.15 |

**Note:** BBB $k_w$ = blood-brain barrier water exchange rate, CBF = cerebral blood flow, eTIV = estimated total intracranial volume. 95% CI = 95 % credible interval**,** *cor* = correlation between the Intercept of region and slope of FW, *sd* = standard deviation of distribution, *sigma* = residual standard deviation.

**eTable 5 – Relationship between BBB $k_w$ and WMH across regions.**

| Parameter | Estimate (Error) | 95% CI (low, upp) | R-hat | ESS | Model Fit |
|---|---|---|---|---|---|
| Intercept | 0.28 (0.18) | -0.07, 0.63 | 1.00 | 914 | |
| $WMH_{log}$ | 0.01 (0.03) | -0.06, 0.07 | 1.00 | 4294 | |
| Age | -0.18 (0.08) | -0.33, -0.03 | 1.00 | 826 | |
| Age^2 | -0.08 (0.06) | -0.20, 0.05 | 1.00 | 844 | |
| Sex (Male) | -0.63 (0.18) | -0.98, -0.28 | 1.00 | 638 | |
| eTIV | 0.11 (0.09) | -0.06, 0.27 | 1.00 | 713 | |
| *sigma* | 0.38 (0.01) | 0.36, 0.40 | 1.00 | 8706 | |
| *sd* of Subject effect (Intercept) | 0..84 (0.05) | 0.74, 0.95 | 1.00 | 1270 | |
| *sd* of Region effect (Intercept) | 0.32 (0.14) | 0.16, 0.66 | 1.00 | 2678 | |
| *sd* of Region effect ($WMH_{log}$ slope) | -0.04 (0.03) | 0.00, 0.12 | 1.00 | 3452 | |
| *cor* of Region effect (Intercept, $WMH_{log}$ slope) | -0.37 (0.40) | -0.93, 0.53 | 1.00 | 6989 | |
| $R^2$ | | | | | 0.86 |
| Adjusted $R^2$ | | | | | 0.82 |
| Marginal $R^2$ | | | | | 0.14 |

**Note:** BBB $k_w$ = blood-brain barrier water exchange rate, $WMH_{log}$ = log transformed white matter hyperintensity volume, eTIV = estimated total intracranial volume. 95% CI = 95 % credible interval**,** *cor* = correlation between the Intercept of region and slope of FW, *sd* = standard deviation of distribution, *sigma* = residual standard deviation.

**eTable 6 – Relationship between Free Water and ePVS across regions.**

| Parameter | Estimate (Error) | 95% CI (low, upp) | R-hat | ESS | Model Fit |
|---|---|---|---|---|---|
| Intercept | 0.46 (0.19) | 0.23, 0.93 | 1.00 | 2118 | |
| ePVS | 0.44 (0.08) | 0.30, 0.63 | 1.00 | 3082 | |
| Age | 0.27 (0.06) | 0.15, 0.40 | 1.00 | 1640 | |
| Age^2 | 0.03 (0.05) | -0.07, 0.13 | 1.00 | 1840 | |
| Sex (Male) | -0.36 (0.15) | -0.66, -0.07 | 1.00 | 1510 | |
| eTIV | 0.18 (0.07) | 0.04, 0.32 | 1.00 | 1575 | |
| *sigma* | 0.51 (0.01) | 0.48, 0.54 | 1.00 | 9986 | |
| *sd* of Subject effect (Intercept) | 0.68 (0.05) | 0.59, 0.78 | 1.00 | 2083 | |
| *sd* of Region effect (Intercept) | 0.46 (0.19) | 0.23, 0.93 | 1.00 | 3272 | |
| *sd* of Region effect (ePVS slope) | 0.10 (0.08) | 0.01, 0.29 | 1.00 | 3296 | |
| *cor* of Region effect (Intercept, ePVS slope) | 0.45 (0.38) | -0.46, 0.95 | 1.00 | 6978 | |
| $R^2$ | | | | | 0.74 |
| Adjusted $R^2$ | | | | | 0.69 |
| Marginal $R^2$ | | | | | 0.26 |

**Note:** FW = Free Water volume fraction, ePVS = enlarged perivascular space volume fraction, eTIV = estimated total intracranial volume. 95% CI = 95 % credible interval**,** *cor* = correlation between the Intercept of region and slope of FW, *sd* = standard deviation of distribution, *sigma* = residual standard deviation.

**eTable 7 – Relationship between Free Water and CBF across regions.**

| Parameter | Estimate (Error) | 95% CI (low, upp) | R-hat | ESS | Model Fit |
|---|---|---|---|---|---|
| Intercept | 0.15 (0.16) | -0.15, 0.45 | 1.00 | 1258 | |
| CBF | 0.07 (0.04) | -0.01, 0.15 | 1.00 | 4458 | |
| Age | 0.24 (0.07) | 0.10, 0.39 | 1.00 | 1362 | |
| Age^2 | 0.02 (0.06) | -0.10, 0.14 | 1.00 | 1439 | |
| Sex (Male) | -0.56 (0.17) | -0.89, -0.23 | 1.00 | 1087 | |
| eTIV | 0.21 (0.08) | 0.05, 0.36 | 1.00 | 1016 | |
| *sigma* | 0.53 (0.01) | 0.50, 0.56 | 1.00 | 9335 | |
| *sd* of Subject effect (Intercept) | 0.79 (0.05) | 0.69, 0.90 | 1.00 | 1981 | |
| *sd* of Region effect (Intercept) | 0.25 (0.12) | 0.12, 0.55 | 1.00 | 2523 | |
| *sd* of Region effect (CBF slope) | 0.03 (0.03) | 0.00, 0.10 | 1.00 | 5661 | |
| *cor* of Region effect (Intercept, CBF slope) | 0.14 (0.45) | -0.76, 0.87 | 1.00 | 112349 | |
| $R^2$ | | | | | 0.73 |
| Adjusted $R^2$ | | | | | 0.67 |
| Marginal $R^2$ | | | | | 0.13 |

**Note:** FW = Free Water volume fraction, CBF = cerebral blood flow, eTIV = estimated total intracranial volume. 95% CI = 95 % credible interval**,** *cor* = correlation between the Intercept of region and slope of FW, *sd* = standard deviation of distribution, *sigma* = residual standard deviation.

**eTable 8 – Relationship between Free Water and WMH across regions.**

| Parameter | Estimate (Error) | 95% CI (low, upp) | R-hat | ESS | Model Fit |
|---|---|---|---|---|---|
| Intercept | 0.15 (0.17) | -0.19, 0.49 | 1.00 | 1659 | |
| $WMH_{log}$ | 0.13 (0.04) | 0.04, 0.21 | 1.00 | 4509 | |
| Age | 0.20 (0.07) | 0.05, 0.35 | 1.00 | 1394 | |
| Age^2 | 0.01 (0.06) | -0.11, 0.12 | 1.00 | 1416 | |
| Sex (Male) | -0.53 (0.17) | -0.87, -0.19 | 1.00 | 1118 | |
| eTIV | 0.19 (0.08) | 0.03, 0.35 | 1.00 | 1069 | |
| *sigma* | 0.52 (0.01) | 0.50, 0.55 | 1.00 | 10094 | |
| *sd* of Subject effect (Intercept) | 0.77 (0.05) | 0.68, 0.88 | 1.00 | 1578 | |
| *sd* of Region effect (Intercept) | 0.30 (0.13) | 0.14, 0.65 | 1.00 | 3070 | |
| *sd* of Region effect ($WMH_{log}$ slope) | 0.04 (0.04) | 0.00, 0.14 | 1.00 | 4121 | |
| *cor* of Region effect (Intercept, $WMH_{log}$ slope) | -0.03 (0.44) | -0.82, 0.78 | 1.00 | 9404 | |
| $R^2$ | | | | | 0.73 |
| Adjusted $R^2$ | | | | | 0.67 |
| Marginal $R^2$ | | | | | 0.13 |

**Note:** FW = Free Water volume fraction, $WMH_{log}$ = log transformed white matter hyperintensity volume, eTIV = estimated total intracranial volume. 95% CI = 95 % credible interval**,** *cor* = correlation between the Intercept of region and slope of FW, *sd* = standard deviation of distribution, *sigma* = residual standard deviation.

**eTable 9 – Relationship between ePVS and CBF across regions.**

| Parameter | Estimate (Error) | 95% CI (low, upp) | R-hat | ESS | Model Fit |
|---|---|---|---|---|---|
| Intercept | 0.19 (0.38) | -0.57, 0.95 | 1.00 | 2144 | |
| CBF | 0.01 (0.04) | -0.06, 0.08 | 1.00 | 4691 | |
| Age | -0.11 (0.05) | -0.21, -0.01 | 1.00 | 1704 | |
| Age^2 | -0.03 (0.04) | -0.11, 0.06 | 1.00 | 1734 | |
| Sex (Male) | -0.49 (0.13) | -0.74, -0.24 | 1.00 | 1441 | |
| eTIV | 0.05 (0.06) | -0.06, 0.17 | 1.00 | 1421 | |
| *sigma* | 0.43 (0.01) | 0.41, 0.45 | 1.00 | 9745 | |
| *sd* of Subject effect (Intercept) | 0.57 (0.04) | 0.50, 0.66 | 1.00 | 1945 | |
| *sd* of Region effect (Intercept) | 0.87 (0.31) | 0.47, 1.68 | 1.00 | 3251 | |
| *sd* of Region effect (CBF slope) | 0.03 (0.02) | 0.00, 0.09 | 1.00 | 5156 | |
| *cor* of Region effect (Intercept, CBF slope) | 0.17 (0.43) | -0.72, 0.88 | 1.00 | 10555 | |
| $R^2$ | | | | | 0.82 |
| Adjusted $R^2$ | | | | | 0.78 |
| Marginal $R^2$ | | | | | 0.07 |

**Note:** ePVS = enlarged perivascular space volume fraction, CBF = cerebral blood flow, eTIV = estimated total intracranial volume. 95% CI = 95 % credible interval**,** *cor* = correlation between the Intercept of region and slope of FW, *sd* = standard deviation of distribution, *sigma* = residual standard deviation.

**eTable 10 – Relationship between ePVS and WMH across regions.**

| Parameter | Estimate (Error) | 95% CI (low, upp) | R-hat | ESS | Model Fit |
|---|---|---|---|---|---|
| Intercept | 0.19 (0.37) | -0.53, 0.96 | 1.00 | 1964 | |
| $WMH_{log}$ | 0.06 (0.04) | -0.03, 0.14 | 1.00 | 5176 | |
| Age | -0.13 (0.05) | -0.24, -0.02 | 1.00 | 1684 | |
| Age^2 | -0.03 (0.04) | -0.12, 0.05 | 1.00 | 1927 | |
| Sex (Male) | -0.48 (0.13) | -0.73, -0.23 | 1.00 | 1538 | |
| eTIV | 0.05 (0.06) | -0.07, 0.16 | 1.00 | 1478 | |
| *sigma* | 0.43 (0.01) | 0.40, 0.45 | 1.00 | 11804 | |
| *sd* of Subject effect (Intercept) | 0.57 (0.04) | 0.50, 0.65 | 1.00 | 2190 | |
| *sd* of Region effect (Intercept) | 0.83 (0.31) | 0.45, 1.59 | 1.00 | 3218 | |
| *sd* of Region effect ($WMH_{log}$ slope) | 0.07 (0.05) | 0.01, 0.18 | 1.00 | 2980 | |
| *cor* of Region effect (Intercept, $WMH_{log}$ slope) | -0.30 (0.37) | -0.87, 0.52 | 1.00 | 8572 | |
| $R^2$ | | | | | 0.82 |
| Adjusted $R^2$ | | | | | 0.78 |
| Marginal $R^2$ | | | | | 0.08 |

**Note:** ePVS = enlarged perivascular space volume fraction, $WMH_{log}$ = log transformed white matter hyperintensity volume, eTIV = estimated total intracranial volume. 95% CI = 95 % credible interval**,** *cor* = correlation between the Intercept of region and slope of FW, *sd* = standard deviation of distribution, *sigma* = residual standard deviation.

**eTable 11 – Relationship between CBF and WMH across regions.**

| Parameter | Estimate (Error) | 95% CI (low, upp) | R-hat | ESS | Model Fit |
|---|---|---|---|---|---|
| Intercept | 0.18 (0.22) | -0.26, 0.62 | 1.00 | 1407 | |
| $WMH_{log}$ | -0.00 (0.04) | -0.07, 0.07 | 1.00 | 4260 | |
| Age | -0.22 (0.08) | -0.37, -0.08 | 1.00 | 922 | |
| Age^2 | -0.10 (0.06) | -0.22, 0.02 | 1.00 | 1116 | |
| Sex (Male) | -0.23 (0.17) | -0.56, 0.10 | 1.00 | 996 | |
| eTIV | 0.07 (0.08) | -0.09, 0.23 | 1.00 | 967 | |
| *sigma* | 0.46 (0.01) | 0.43, 0.48 | 1.00 | 10026 | |
| *sd* of Subject effect (Intercept) | 0.80 (0.05) | 0.71, 0.91 | 1.00 | 1515 | |
| *sd* of Region effect (Intercept) | 0.44 (0.18) | 0.22, 0.90 | 1.00 | 2885 | |
| *sd* of Region effect ($WMH_{log}$ slope) | 0.03 (0.03) | 0.00, 0.10 | 1.00 | 5105 | |
| *cor* of Region effect (Intercept, $WMH_{log}$ slope) | -0.08 (0.44) | -0.84, 0.77 | 1.00 | 9660 | |
| $R^2$ | | | | | 0.79 |
| Adjusted $R^2$ | | | | | 0.75 |
| Marginal $R^2$ | | | | | 0.10 |

**Note:** CBF = cerebral blood flow, $WMH_{log}$ = log transformed white matter hyperintensity volume, eTIV = estimated total intracranial volume. 95% CI = 95 % credible interval**,** *cor* = correlation between the Intercept of region and slope of FW, *sd* = standard deviation of distribution, *sigma* = residual standard deviation.

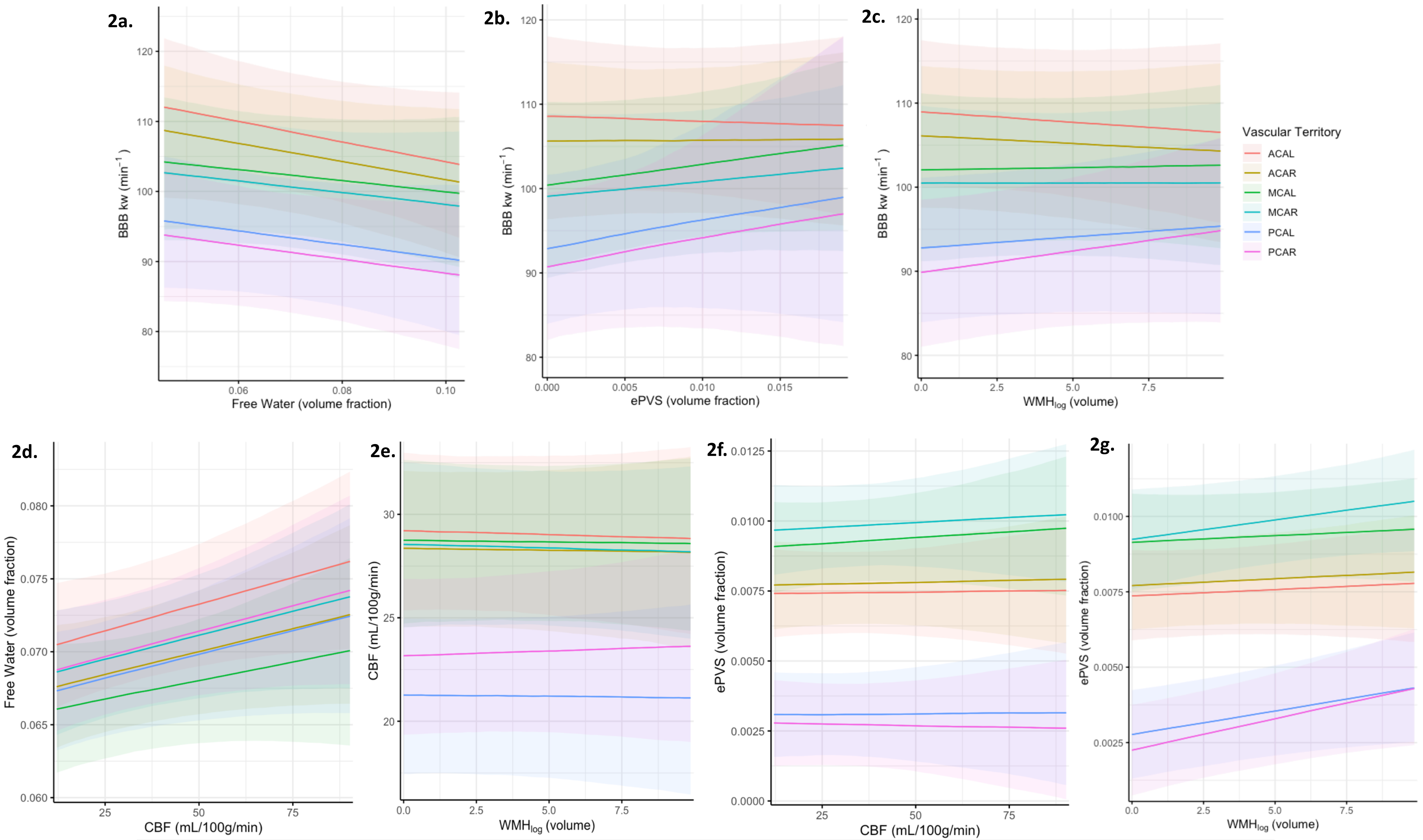


**eFigure2 – Plotted relationships between MRI measure pairs.**

a) BBB $k_w$ and Free Water; b) BBB $k_w$ and ePVS; c) BBB $k_w$ and WMH; d) Free Water and CBF; e) CBF and WMH; f) ePVS and CBF; g) ePVS and WMH. BBB $k_w$ = blood-brain barrier water exchange rate; CBF = gray matter cerebral blood flow; ePVS = enlarged perivascular space volume as a fraction of total normal appearing white matter and gray matter; FW = Free Water volume fraction; $WMH_{log}$ = log transformed white matter hyperintensity volume.

**eTable 12 – Relationship between MRI measures and Age.**

| Model | $\beta$ (SE) | t-value | p-value | $R^2_{Adjusted}$ |
|---|---|---|---|---|
| BBB kw | | | | 0.123 |
| Intercept | 0.216 (0.103) | 2.095 | .038 | |
| Age (total) | - | - | .017 | |
| Age | -2.459 (0.949) | 2.590 | .011 | |
| $Age^2$ | -1.192 (0.943) | 1.263 | .209 | |
| Sex (Male) | -0.678 (0.198) | 3.428 | <.001 | |
| eTIV | 0.133 (0.092) | 1.439 | .153 | |
| FW | | | | 0.132 |
| Intercept | 0.217 (0.102) | 2.121 | .036 | |
| Age (total) | - | - | .004 | |
| Age | 3.111 (0.944) | 3.295 | .001 | |
| $Age^2$ | 0.596 (0.938) | 0.636 | .526 | |
| Sex (Male) | -0.683 (0.197) | 3.470 | <.001 | |
| eTIV | 0.256 (0.092) | 2.778 | .006 | |
| ePVS | | | | 0.136 |
| Intercept | 0.239 (0.102) | 2.340 | .021 | |
| Age (total) | - | - | .049 | |
| Age | -2.238 (0.942) | 2.375 | .019 | |
| $Age^2$ | -0.629 (0.936) | 0.672 | .503 | |
| Sex (Male) | -0.752 (0.196) | 3.828 | <.001 | |
| eTIV | 0.075 (0.092) | 0.814 | .417 | |
| CBF | | | | 0.213 |
| Intercept | 0.103 (0.098) | 1.057 | .292 | |
| Age (total) | - | - | <.001 | |
| Age | -4.579 (0.899) | 5.095 | <.001 | |
| $Age^2$ | -0.119 (0.893) | 0.133 | .894 | |
| Sex (Male) | -0.324 (0.187) | 1.730 | .086 | |
| eTIV | -0.107 (0.088) | 1.227 | .222 | |
| $WMH_{log}$ | | | | 0.126 |
| Intercept | 0.127 (0.103) | 1.235 | .219 | |
| Age (total) | - | - | <.001 | |
| Age | 3.632 (0.948) | 3.833 | <.001 | |
| $Age^2$ | 0.649 (0.942) | 0.689 | .492 | |
| Sex (Male) | -0.399 (0.198) | 2.020 | .045 | |
| eTIV | 0.238 (0.092) | 2.574 | .011 | |

**Note:** $\beta$ = standardised estimate; SE = standard error.
BBB $k_w$ = blood-brain barrier water exchange rate; FW = Free Water volume fraction;
ePVS = enlarged perivascular space volume fraction; CBF = gray matter cerebral blood flow;
$WMH_{log}$ = log transformed white matter hyperintensity volume.

**eTable 13 – Age interactions on relationships between MRI pairs.**

| Model | $\beta$ (SE) | t-value | p-value | $R^2_{Adjusted}$ |
|---|---|---|---|---|
| FW x BBB kw | | | | 0.127 |
| Intercept | 0.192 (0.105) | 1.824 | .071 | |
| BBB kw | 0.056 (0.091) | 0.613 | .541 | |
| Age (total) | | | .005 | |
| Age | 2.964 (1.029) | 2.881 | .005 | |
| $Age^2$ | 0.053 (1.036) | 0.051 | .959 | |
| Sex (Male) | -0.691 (0.209) | 3.309 | .001 | |
| eTIV | 0.259 (0.093) | 2.780 | .006 | |
| BBB kw*Age (total) | | | .354 | |
| BBB kw*Age | -1.209 (0.892) | 1.356 | .178 | |
| BBB kw*$Age^2$ | -0.192 (0.793) | 0.242 | .809 | |
| ePVS x BBB kw | | | | 0.124 |
| Intercept | 0.232 (0.106) | 2.201 | .030 | |
| BBB kw | -0.010 (0.091) | 0.115 | .909 | |
| Age (total) | | | .054 | |
| Age | -2.418 (1.031) | 2.346 | .021 | |
| $Age^2$ | -1.106 (1.038) | 1.066 | .289 | |
| Sex (Male) | -0.795 (0.209) | 3.797 | .000 | |
| eTIV | 0.084 (0.093) | 0.900 | .370 | |
| BBB kw*Age (total) | | | .561 | |
| BBB kw*Age | -0.943 (0.893) | 1.056 | .293 | |
| BBB kw*$Age^2$ | -0.013 (0.794) | 0.017 | .987 | |
| CBF x BBB kw | | | | 0.226 |
| Intercept | 0.113 (0.099) | 1.134 | .259 | |
| BBB kw | -0.004 (0.086) | 0.044 | .965 | |
| Age (total) | | | <.001 | |
| Age | -4.765 (0.968) | 4.921 | <.001 | |
| $Age^2$ | 0.662 (0.975) | 0.679 | .499 | |
| Sex (Male) | -0.260 (0.197) | 1.321 | .189 | |
| eTIV | -0.118 (0.088) | 1.340 | .183 | |
| BBB kw*Age (total) | | | .083 | |
| BBB kw*Age | 1.755 (0.839) | 2.091 | .039 | |
| BBB kw*$Age^2$ | -0.914 (0.746) | 1.226 | .223 | |
| $WMH_{log}$ x BBB kw | | | | 0.130 |
| Intercept | 0.122 (0.105) | 1.159 | .249 | |
| BBB kw | -0.041 (0.091) | 0.446 | .656 | |
| Age (total) | | | .002 | |
| Age | 3.357 (1.027) | 3.270 | .001 | |
| $Age^2$ | -0.175 (1.034) | 0.170 | .866 | |
| Sex (Male) | -0.488 (0.208) | 2.340 | .021 | |
| eTIV | 0.256 (0.093) | 2.750 | .007 | |
| BBB kw*Age (total) | | | .199 | |
| BBB kw*Age | -1.605 (0.890) | 1.804 | .074 | |
| BBB kw*$Age^2$ | 0.154 (0.791) | 0.195 | .845 | |

**eTable 13. continued**

| Model | $\beta$ (SE) | t-value | p-value | $R^2_{Adjusted}$ |
|---|---|---|---|---|
| ePVS x FW | | | | 0.378 |
| Intercept | 0.129 (0.092) | 1.407 | .162 | |
| FW | 0.543 (0.076) | 7.175 | <.001 | |
| Age (total) | | | <.001 | |
| Age | -3.996 (0.867) | 4.611 | <.001 | |
| $Age^2$ | -0.953 (0.853) | 1.118 | .266 | |
| Sex (Male) | -0.394 (0.177) | 2.231 | .028 | |
| eTIV | -0.064 (0.080) | 0.793 | .429 | |
| FW*Age (total) | | | .879 | |
| FW*Age | -0.237 (0.825) | 0.287 | .774 | |
| FW*$Age^2$ | 0.308 (0.703) | 0.438 | .662 | |
| CBF x FW | | | | 0.231 |
| Intercept | 0.045 (0.102) | 0.440 | .661 | |
| FW | -0.030 (0.084) | 0.358 | .721 | |
| Age (total) | | | <.001 | |
| Age | -4.390 (0.963) | 4.557 | <.001 | |
| $Age^2$ | 0.429 (0.948) | 0.453 | .651 | |
| Sex (Male) | -0.256 (0.197) | 1.305 | .194 | |
| eTIV | -0.099 (0.089) | 1.106 | .271 | |
| FW*Age (total) | | | .059 | |
| FW*Age | 1.868 (0.917) | 2.037 | .044 | |
| FW*$Age^2$ | -1.116 (0.782) | 1.427 | .156 | |
| $WMH_{log}$ x FW | | | | 0.132 |
| Intercept | 0.070 (0.109) | 0.645 | .520 | |
| FW | 0.143 (0.089) | 1.598 | .112 | |
| Age (total) | | | .006 | |
| Age | 3.275 (1.024) | 3.199 | .002 | |
| $Age^2$ | 0.471 (1.007) | 0.468 | .641 | |
| Sex (Male) | -0.264 (0.209) | -1.264 | .208 | |
| eTIV | 0.201 (0.095) | 2.116 | .036 | |
| FW*Age (total) | | | .606 | |
| FW*Age | 0.738 (0.974) | 0.758 | .450 | |
| FW*$Age^2$ | -0.589 (0.831) | -0.709 | .480 | |
| CBF x ePVS | | | | 0.201 |
| Intercept | 0.115 (0.102) | 1.130 | .261 | |
| ePVS | -0.080 (0.086) | 0.934 | .352 | |
| Age (total) | | | <.001 | |
| Age | -4.833 (0.943) | 5.126 | <.001 | |
| $Age^2$ | -0.240 (0.917) | 0.262 | .794 | |
| Sex (Male) | -0.373 (0.201) | 1.856 | .066 | |
| eTIV | -0.099 (0.089) | 1.116 | .267 | |
| ePVS*Age (total) | | | .889 | |
| ePVS*Age | -0.077 (1.029) | 0.075 | .940 | |
| ePVS*$Age^2$ | 0.487 (1.071) | 0.455 | .650 | |

**eTable 13. continued**

| Model | $\beta$ (SE) | t-value | p-value | $R^2_{Adjusted}$ |
|---|---|---|---|---|
| $WMH_{log}$ x ePVS | | | | 0.158 |
| Intercept | 0.069 (0.105) | 0.655 | .514 | |
| ePVS | 0.042 (0.088) | 0.478 | .634 | |
| Age (total) | | | <.001 | |
| Age | 3.289 (0.968) | 3.398 | <.001 | |
| $Age^2$ | 0.239 (0.941) | 0.254 | .800 | |
| Sex (Male) | -0.321 (0.206) | 1.557 | .122 | |
| eTIV | 0.246 (0.091) | 2.701 | .008 | |
| ePVS*Age (total) | | | .029 | |
| ePVS*Age | -1.220 (1.057) | 1.154 | .251 | |
| ePVS*$Age^2$ | -2.364 (1.100) | 2.150 | .034 | |
| $WMH_{log}$ x CBF | | | | 0.116 |
| Intercept | 0.085 (0.111) | 0.763 | .447 | |
| CBF | 0.065 (0.095) | 0.688 | .492 | |
| Age (total) | | | <.001 | |
| Age | 3.540 (1.151) | 3.077 | .003 | |
| $Age^2$ | 0.147 (1.080) | 0.136 | .892 | |
| Sex (Male) | -0.383 (0.201) | 1.904 | .059 | |
| eTIV | 0.258 (0.094) | 2.732 | .007 | |
| CBF*Age (total) | | | .613 | |
| CBF*Age | -1.005 (1.066) | 0.944 | .347 | |
| CBF*$Age^2$ | -0.634 (0.991) | 0.639 | .524 | |

**Note:** $\beta$ = standardised estimate; SE = standard error.
BBB $k_w$ = blood-brain barrier water exchange rate; FW = Free Water volume fraction;
ePVS = enlarged perivascular space volume fraction; CBF = gray matter cerebral blood flow;
$WMH_{log}$ = log transformed white matter hyperintensity volume.

**eTable 14 – Relationship between MRI measures and brain age.**

| Model | $\beta$ (SE) | t-value | p-value | $R^2_{Adjusted}$ |
|---|---|---|---|---|
| BBB kw | | | | 0.118 |
| Intercept | 0.214 (0.103) | 2.067 | .041 | |
| brainPAD | -0.047 (0.833) | 0.566 | .572 | |
| Age (total) | - | - | .015 | |
| Age | -2.486 (0.953) | 2.609 | .010 | |
| $Age^2$ | -1.260 (0.953) | 1.321 | .189 | |
| Sex (Male) | -0.671 (0.199) | 3.378 | <.001 | |
| eTIV | 0.136 (0.093) | 1.461 | .147 | |
| FW | | | | 0.127 |
| Intercept | 0.220 (0.103) | 2.137 | .035 | |
| brainPAD | 0.049 (0.082) | 0.593 | .554 | |
| Age (total) | - | - | .004 | |
| Age | 3.140 (0.948) | 3.312 | .001 | |
| $Age^2$ | 0.667 (0.948) | 0.704 | .483 | |
| Sex (Male) | -0.690 (0.198) | 3.491 | <.001 | |
| eTIV | 0.253 (0.092) | 2.740 | .007 | |
| ePVS | | | | 0.150 |
| Intercept | 0.232 (0.101) | 2.291 | .024 | |
| brainPAD | -0.144 (0.082) | 1.757 | .081 | |
| Age (total) | - | - | .033 | |
| Age | -2.321 (0.936) | 2.480 | .014 | |
| $Age^2$ | -0.836 (0.936) | 0.894 | .373 | |
| Sex (Male) | -0.731 (0.195) | 3.744 | <.001 | |
| eTIV | 0.082 (0.091) | 0.903 | .368 | |
| CBF | | | | 0.221 |
| Intercept | 0.098 (0.097) | 1.006 | .316 | |
| brainPAD | -0.115 (0.078) | 1.471 | .144 | |
| Age (total) | - | - | <.001 | |
| Age | -4.646 (0.896) | 5.186 | <.001 | |
| $Age^2$ | -0.285 (0.896) | 0.318 | .751 | |
| Sex (Male) | -0.307 (0.187) | 1.644 | .103 | |
| eTIV | -0.101 (0.087) | 1.162 | .248 | |
| $WMH_{log}$ | | | | 0.119 |
| Intercept | 0.127 (0.103) | 1.228 | .222 | |
| brainPAD | -0.003 (0.083) | 0.037 | .970 | |
| Age (total) | - | - | <.001 | |
| Age | 3.631 (0.953) | 3.812 | <.001 | |
| $Age^2$ | 0.644 (0.953) | 0.676 | .500 | |
| Sex (Male) | -0.399 (0.199) | 2.006 | .047 | |
| eTIV | 0.238 (0.093) | 2.563 | .012 | |

**Note:** $\beta$ = standardised estimate; SE = standard error.
BBB $k_w$ = blood-brain barrier water exchange rate; FW = Free Water volume fraction;
ePVS = enlarged perivascular space volume fraction; CBF = gray matter cerebral blood flow;
$WMH_{log}$ = log transformed white matter hyperintensity volume.

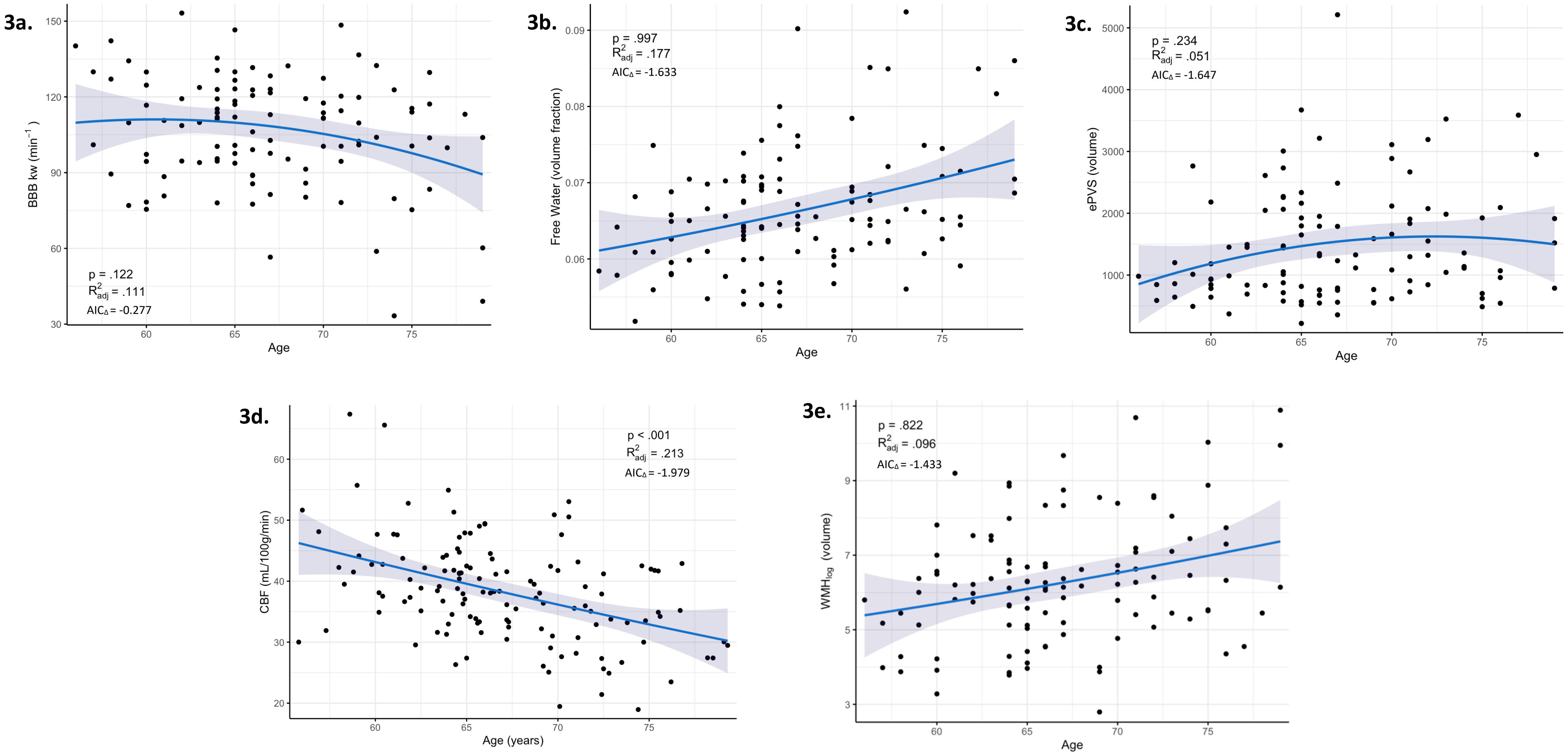


**eFigure 3.** Associations between MRI outcomes and age with a quadratic fit.

a) BBB $k_w$ and age; b) FW and age; c) ePVS and age; d) CBF and age; e) WMH and age.
BBB $k_w$ = blood-brain barrier water exchange rate; CBF = gray matter cerebral blood flow;
ePVS = enlarged perivascular space volume fraction; FW = Free Water volume fraction; $WMH_{log}$ = log transformed white matter hyperintensity volume.
$AIC_\Delta$ values represent the linear age model AIC minus the quadratic age model AIC (negative values favour linear model).
